# Polygenic Scores for Cognitive Abilities and their Association with Different Aspects of General Intelligence – a Deep Phenotyping Approach

**DOI:** 10.1101/2020.06.03.131318

**Authors:** Erhan Genç, Caroline Schlüter, Christoph Fraenz, Larissa Arning, Dorothea Metzen, Huu Phuc Nguyen, Manuel C. Voelkle, Fabian Streit, Onur Güntürkün, Robert Kumsta, Sebastian Ocklenburg

## Abstract

Intelligence is a highly polygenic trait and genome-wide association studies (GWAS) have identified thousands of DNA variants contributing with small effects. Polygenic scores (PGS) can aggregate those effects for trait prediction in independent samples. As large-scale light-phenotyping GWAS operationalized intelligence as performance in rather superficial tests, the question arises which intelligence facets are actually captured. We used deep-phenotyping to investigate the molecular determinants of individual differences in cognitive ability. We therefore studied the association between PGS of intelligence (IQ-PGS), cognitive performance (CP-PGS) and educational attainment (EA-PGS) with a wide range of intelligence facets in a sample of 557 healthy adults. IQ-PGS, CP-PGS and EA-PGS had the highest incremental *R*^2^s for general (2.71%; 4.27%; 2.06%), verbal (3.30%; 4.64%; 1.61%) and numerical intelligence (3.06%; 3.24%; 1.26%) and the weakest for non-verbal intelligence (0.89%; 1.47%; 0.70%) and memory (0.80%; 1.06%; 0.67%). These results indicate that PGS derived from light-phenotyping GWAS do not reflect different facets of intelligence equally well, and thus should not be interpreted as genetic indicators of intelligence per se. The findings refine our understanding of how PGS are related to other traits or life outcomes.

## Introduction

Gaining insight into the molecular determinants of differences in cognitive abilities is one of the core aims of neurobiological intelligence research. Our ability to “understand complex ideas, to adapt effectively to the environment, to learn from experience, to engage in various forms of reasoning [and] to overcome obstacles by taking thought” has usually been described as general intelligence [1]. Various tests have been designed to measure the cognitive abilities of a person by assessing different aspects of general intelligence, like inductive and deductive reasoning abilities or the amount of acquired declarative knowledge [2]. Whereas most test procedures cover different aspects of general intelligence, there are also tests that focus on specific cognitive abilities. Matrix reasoning tests, for instance, are typically used to assess non-verbal abstract reasoning [3,4], while other tests measure stored long-term memory of static information like rules, relationships, abstract concepts, and of course, general knowledge [2,5].

Decades of intelligence research have shown that general intelligence is one of the best predictors of important life outcomes, including educational and occupational success [6,7] as well as mental and physical health [8-10]. Thus, considerable research efforts have been put into place to explore the mechanisms behind interindividual differences in general intelligence. Behavioral genetics has been particularly fruitful for intelligence research. Twin and family studies have demonstrated that general intelligence is one of the most heritable behavioral traits, with heritability estimates ranging from 60% to 80% in adulthood [for an overview see 11,12]. However, these traditional quantitative genetics studies cannot be used to estimate which and how many genetic variants contribute to heritability. Although, as early as 1918, Fisher’s infinitesimal model postulated that complex traits are affected by a large number of genes, it was not until the advent of genome-wide association studies (GWAS) that effect sizes of single nucleotide polymorphisms (SNPs) could be systematically assessed over the genome. In support of Fisher’s infinitesimal model [13], large GWAS on intelligence have shown that even the most strongly associated SNPs explain less than 1% of the variance, and that heritability of intelligence is caused by a very large number of DNA variants of small effect (not taking into account rare mutations with potentially large effects on individuals, but small effects on the population) [14].

Given that GWAS of sufficient sample size can reliably detect very small effects of single common variants, and given that SNPs contribute cumulatively to heritability, a fruitful approach forward has been the use of so-called polygenic scores (PGS). These are genetic indices of a trait, defined as the sum of trait-associated alleles across many genetic loci, weighted by effect sizes estimated by GWAS. Such scores can be calculated for individuals in target samples (independent from the initial discovery GWAS) and be used to predict traits of interest [15]. For instance, PGS for intelligence (IQ-PGS) [16,17] and cognitive performance (CP-PGS) as well as educational attainment (EA-PGS) [18-20], a secondary measure of intelligence, have been associated with a wide variety of traits, including life-course development, educational achievement, body mass index or emotional and behavioral problems in children [21-23]. Although IQ-PGS, CP-PGS and EA-PGS explain a considerable amount of variance in intelligence (which is thought to increase even further with larger GWAS) [14], and robust and sometimes unexpected associations between genetic indices of cognitive abilities and other traits have been uncovered, it is important to understand that the predictive power of these PGS depends on the cognitive measure that is being used. To reliably identify genetic variants associated with a complex continuous behavioral trait, such as intelligence, in a GWAS, large samples sizes in the 100,000s to millions are required. This has been successfully achieved using a light-phenotyping approach, that is, performing GWAS on the performance in rather superficial tests of general cognitive abilities [24,25], or even more crudely, years of education [25]. The question thus arises, which of the various aspects of general intelligence [2,1], are mainly reflected in those GWAS. The study at hand aimed to tackle this issue by pursuing a deep phenotyping approach. Using an extensive test battery comprised of tests for memory performance, processing speed, reasoning, and general knowledge, we investigated the predictive power of IQ-PGS [24], CP-PGS and EA-PGS [25] with regard to each of the aforementioned cognitive abilities.

## Methods

### Sample size estimation

A statistical power analysis was performed for sample size estimation based on data reported by Savage et al. [24] and Lee et al. [25]. Both meta-analyses reported effect sizes [26] of *R*^2^ = .052 (IQ-PSG) [24], and *R*2 =.097 (EA-PGS) [25], respectively. G-power [27] was used to determine the sample size required to detect a small to medium effect size (*f*^2^ = .08) in a multiple linear regression analysis, using an α error of .05 and a statistical power of 1-β = .90. Sample size estimation predicted that *N* = 236 participants were needed to obtain the desired statistical power. With a final sample size of *N* = 518 (see below), our sample was thus adequately powered for the main objective of this study.

### Participants

We investigated 557 neurologically and psychologically healthy participants with a mean age of 27.33 years (*SD* = 9.43; range 18 - 75 years), including 283 males (mean age: 27.71 years, *SD* = 9.86 years) and 274 females (mean age: 26.94 years, *SD* = 8.96 years). The sample was mainly comprised of university students of different majors (mean years of education: 17.14 years, *SD* = 3.12 years), who either received a financial reward or course credits for their participation. Health status was self-reported by the participants as part of the demographic questionnaire. Individuals who reported current or past neurological or psychological problems were not admitted to the study. The study protocol was approved by the local ethics committee of the Faculty of Psychology at Ruhr University Bochum (vote Nr. 165). All participants gave written informed consent and were treated following the Declaration of Helsinki.

### Acquisition and Analysis of Behavioral Data

Behavioral data was acquired during four individual test sessions. Each session was designed as a group setting of up to six participants, seated at individual tables, in a quiet and well-lit room. The tests were administered according to their respective manuals. The following is a brief description of each test procedure used in our study. Please refer to the Supplementary Material for descriptive statistics (Supplementary Material S1) and intercorrelations (Supplementary Material S2) of all cognitive tests.

#### I-S-T 2000 R

The *Intelligenz-Struktur-Test 2000 R* (I-S-T 2000 R) is a well-established German intelligence test battery measuring multiple facets of general intelligence [28,29]. The test consists of a basic and an extension module. The basic module measures different aspects of intelligence and contains 180 items assessing verbal, numerical and figural abilities as well as 23 items assessing verbal and figural memory. Verbal, numeric, and figural abilities are measured by three reasoning tasks that comprise 20 items each. For instance, verbal intelligence is assed via items on sentences completion, where the participant is asked to complete a sentence with the correct word, or on analogies and commonalities. Numerical intelligence on the contrary comprises items on arithmetic problems, digit spans, and arithmetic operators assessing the mathematical abilities of the participant. Figural intelligence is assessed via items testing for the participant’s ability to assemble figures mentally, to mentally rotate and match dices, and to solve matrix-reasoning tasks. The processing time for this section is about 90 minutes. Subsequently, the participants complete two memory tasks, one verbal (10 items) and one figural (13 items), where they must memorize a series of words, or pairs of figures, respectively. This takes about 10 minutes. The extension module measures general knowledge covering a total number of 84 items. The knowledge test covers verbal (26 questions), numerical (25 questions), and figural knowledge (22 questions) and takes about 40 minutes. Here, the participant’s knowledge on various facets is assessed: Art/literature, geography/history, mathematics, science, and daily life. Most of the items of both modules are designed in multiple-choice form. The only exception are two sub-tests on numerical reasoning (calculations and number series). Here the participant has to directly fill-in the answer. The complete testing session takes about 2 h 30 min. The reliability estimates (Cronbach’s α) for the sub-factettes of the basic module fall between .88 and .95, as well as .93 for the memory tasks and .96 for general mental ability. The extension module has a reliability of .93 (Cronbach’s α). The recent norming sample consists of about 5800 individuals for the basic module and 661 individuals for the extension module. The age range in the norming sample is between 15 - 60 years and both sexes are represented equally.

#### ZVT

The *Zahlenverbindungstest* (ZVT) is a trail-making test used to assess processing speed in both children and adults [30]. After editing two sample matrices, the participant has to process a total of four matrices. Here, the individual has to connect circled numbers from 1 to 90 in ascending order. The numbers were positioned more or less randomly within the matrix. The instructor measures the processing time for each matrix. The total test value, reflecting the participant’s processing speed, is then calculated as the arithmetic mean of all four matrices. The test takes about 10 minutes in total. The reliability between the individual matrices is above .86 and .95 for adults and six months retest-reliability is between .84 and .90. The recent norming sample consists of about 2109 individuals with an age range between 8 - 60 years and equal sex representation.

#### BOMAT advanced short

The *Bochumer Matrizentest* (BOMAT) is a non-verbal intelligence test which is widely used in neuroscientific research [31-33,3]. Its structure is similar to the well-established Raven’s Advanced Progressive Matrices [4]. Within the framework of our study, we carried out the advanced short version of the BOMAT, which is known to have a high discriminatory power in samples with generally high intellectual abilities, thus avoiding possible ceiling effects [33,32]. The test comprises two parallel forms with 29 matrix-reasoning items each. Participants were assigned to one of the two forms, which they had to complete. The participants have a total of 45 minutes to process as many matrices as possible. The split-half reliability of the BOMAT is .89, Cronbach’s α is .92, and reliability between the parallel-forms is .86. The recent norming sample consists of about 2100 individuals with an age range between 18 - 60 years and equal sex representation.

#### BOWIT

The *Bochumer Wissenstest* (BOWIT) is a German inventory to assess the subject’s degree of general knowledge [5]. The inventory comprises two parallel forms with 154 items each. Both forms include eleven different facets of general knowledge: Arts/architecture, language/ literature, geography/logistics, philosophy/religion, history/archeology, economics/law, civics/ politics, as well as, biology/chemistry, nutrition/health, mathematics/physics, and technology/ electronics. Within one test form, each knowledge facet is represented by 14 multiple-choice items. In our study, all participants had to complete both test forms resulting in a total number of 308 items. The processing time for each test form is about 45 minutes. The knowledge facets assessed by the BOWIT are very similar to general knowledge inventories used in other studies [34-36]. The inventory fulfills all important quality criteria regarding different measures of reliability and validity. The inventory’s manual specifies that split-half reliability is .96, Cronbach’s α is .95, test-retest reliability is .96 and parallel-form reliability is .91. Convergent and discriminant validity are given for both test forms. The norming sample consists of about 2300 individuals (age range: 18 - 66 years) and has an equal sex representation.

### DNA sampling and genotyping

For non-invasive sampling, exfoliated cells were brushed from the oral mucosa of the participants. DNA isolation was performed with QIAamp DNA mini Kit (Qiagen GmbH, Hilden, Germany). Genotyping was carried out using the Illumina Infinium Global Screening Array 1.0 with MDD and Psych content (Illumina, San Diego, CA, USA) at the Life & Brain facilities, Bonn, Germany. Filtering was performed with PLINK 1.9 [37,38] removing SNPs with a minor allele frequency of <0.01, deviating from Hardy-Weinberg equilibrium with a *p*-value of <1*10-6, and missing data >0.02. Participants were excluded with >0.02 missingness, sex-mismatch, and heterozygosity rate > |0.2|. Filtering for relatedness and population structure was carried out on a SNP set of filtered for high quality (HWE *p* > 0.02, MAF > 0.2, missingness = 0), and LD pruning (*r*^2^ = 0.1). In pairs of cryptically related subjects pi hat > 0.2 was applied to excluded subjects at random. Principal components to control for population stratification were generated, and outliers > |6SD| on one of the first 20 PC were excluded. The final data set consisted of 518 participants and 494,740 SNPs.

### Polygenic scores

We created genome-wide polygenic scores for each participant using publicly available summary statistics for intelligence (N = 269,867), cognitive performance (N = 257,828) and educational attainment (excl. 23andMe; N = 766,345) [25,24]. Polygenic scores were constructed as the weighted sums of each participant’s trait-associated alleles across all SNPs using PRSice 2.1.6 [39]. In regard to the highly polygenic nature identified for EA and IQ and the observed range of highest prediction of the PGS in the original manuscripts [24,25], we applied a *p*-value threshold (PT) of 0.05 for the inclusion of SNPs in the calculation of IQ-PGS, CP-PGS and EA-PGS. Additionally, we report the results for the PGS with the strongest association with the respective cognitive tests (and subtests) in our sample (best-fit PGS). That is, the *p*-value threshold (PT) for inclusion of SNPs was chosen empirically (for the range of PT 5*10-8 - .5 in steps of 5*10-5), so the resulting PGS explained the maximum amount of test score variance for the respective measure in our sample. Finally, we also investigated the predictive power of PGS including all available SNPs (non-fit PGS), that is, the *p*-value threshold for SNP inclusion equaled PT = 1.00. The predictive power of the PGS derived from the GWAS was measured by the ‘incremental *R*^2^’ statistic [25]. The incremental *R*^2^ reflects the increase in the determination coefficient (*R*^2^) when the PGS is added to a regression model predicting the behavioral phenotype alongside a number of control variables (here: sex, age, and the first four principal components of population stratification). For all statistical analyses in PRSice, linear parametric methods were used. Testing was two-tailed with an α-level of *p* < .05. As we report a total of 81 regression analyses, our results were FDR corrected for multiple comparisons and the corrected α-levels were in the range between .05 / 81 = .000617 and .05 as defined by the Benjamini-Hochberg method [40]. Since the sexes differed significantly for some of the phenotypes (see results), we also calculated the above-mentioned analyses in an exploratory fashion separately for the sexes. Control variables were age, and the first four principal components of population stratification. For PT = 0.05 and PT = 1 we applied the same procedure as above. For the best-fit approach, we chose the full sample best-fit PT of the respective phenotype and applied it to the subsamples. Finally, we test whether the predictive power of PGS differentiate between females and males by comparing the incremental *R*^2^ between the two groups correcting for multiple comparisons using the Benjamini-Hochberg method as described above.

## Results

In the following section, we report incremental determination coefficients (incremental *R*^2^) for the PGS with a PT of 0.05 and respective test scores. Additionally, we investigated the association between our cognitive test scores and, so-called best-fit IQ-PGS, CP-PGS and EA-PGS. These best-fit PGS were estimated by using a function, which empirically determines a *p*-value threshold for SNP inclusion to maximally predict the performance in the respective cognitive test (see Supplementary Material S3, S4 and S5 for intercorrelations between best-fit PGS). Finally, we explored the predictive power of non-fit PGS including all available SNPs (PT = 1.00) and cognitive test scores.

For PT of 0.05 IQ-PGS were especially predictive of individual differences in verbal (incremental *R*^2^ = 3.29%, *p* < .001) and numerical intelligence (incremental *R*^2^ = 3.05%, *p* < .001) measured with the IST-2000-R. The predictive power for differences in general intelligence was only slightly lower (Figure 1). Here, IQ-PGS had an incremental *R*^2^ of 2.71% (*p* < .001). Regarding measures of general knowledge, IQ-PGS had an incremental *R*^2^ of 1.85% for ability differences in the IST-2000-R general knowledge test (*p* < .001) and 1.34% for differences assessed with the BOWIT (*p* = .001). Differences in non-verbal aspects of intelligence were weaker predicted. Our multiple regression analyses resulted in predictive values of *R*^2^ = 1.04% for processing speed (*p* = .017), *R*^2^ = 0.89% for matrices (*p* = .029) and *R*^2^ = 0.13% for figural intelligence (*p* = .406). Also, the predictive power of IQ-PGS for memory was weak (incremental *R*^2^ = 0.80%, *p* = .033).

**Figure 1.**
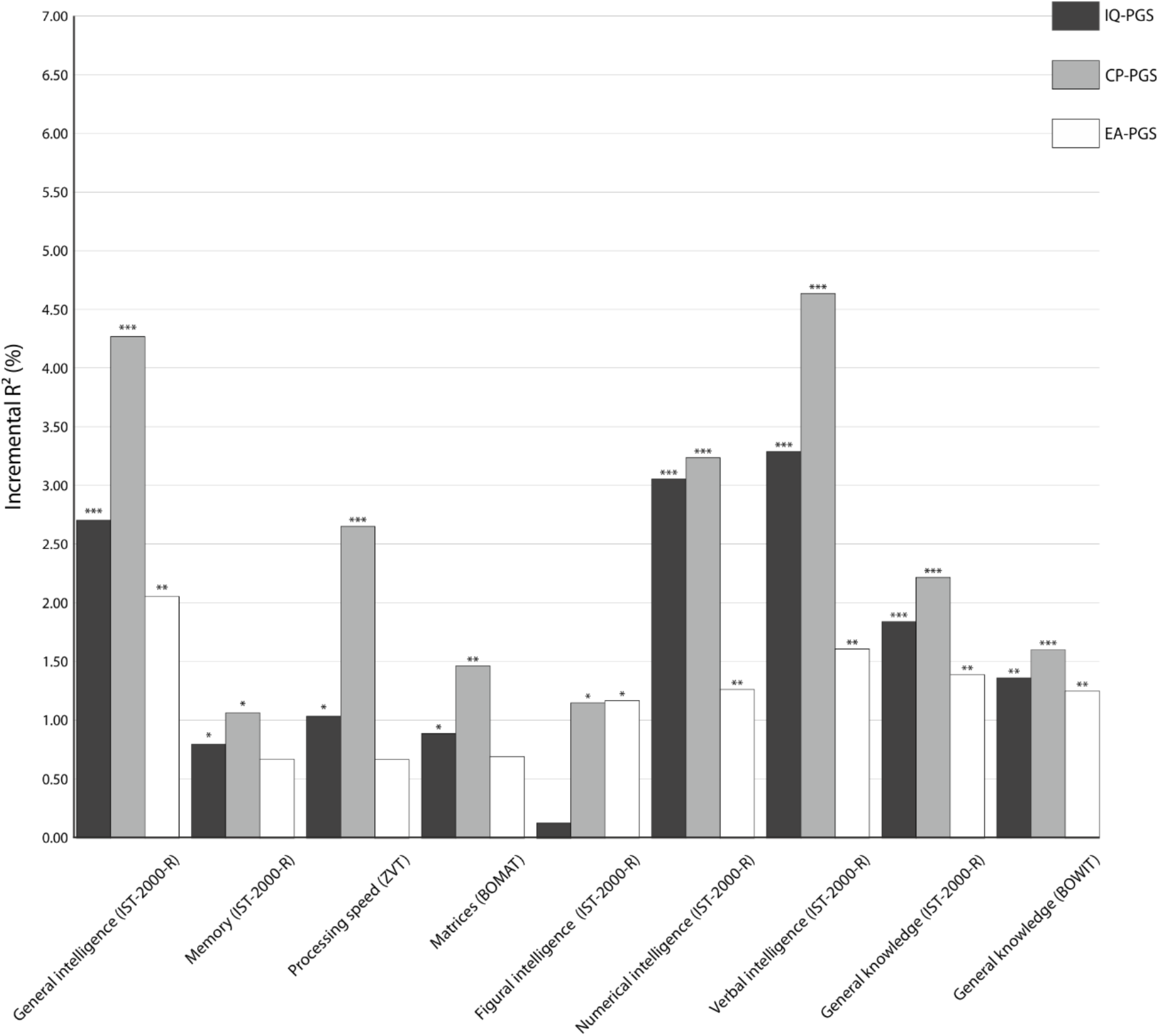
Incremental *R*^2^ of the *P*-value threshold (PT) = .05 polygenic scores of intelligence (IQ-PGS), cognitive performance (CP-PGS) and educational attainment (EA-PGS) in percent. The incremental *R*^2^ reflects the increase in the determination coefficient (*R*^2^) when the IQ-PGS or CP-PGS or EA-PGS is added to a regression model predicting individual differences in the respective cognitive test. The association between PGS and phenotype was controlled for the effects of sex, age, population stratification, and multiple comparisons [40]. * adjusted *p* ≤ .05, ** adjusted *p* ≤ .01, *** adjusted *p* ≤ .001.

Individual differences in CP-PGS were especially predictive of individual differences in verbal (incremental *R*^2^ = 4.64%, *p* < .001) and general intelligence (incremental *R*^2^ = 4.27%, *p* < .0021) measured with the IST-2000-R. The predictive power for differences in numerical intelligence was only slightly lower (Figure 1). Here, CP-PGS had an incremental *R*^2^ of 3.24% (*p* < .001). Regarding measures of general knowledge, CP-PGS had an incremental *R*^2^ of 2.22% for ability differences in the IST-2000-R general knowledge test (*p* < .001) and 1.60% for differences assessed with the BOWIT (*p* < .001). Differences in language-free components of general intelligence, measured as figural intelligence (incremental *R*^2^ = 1.15%, *p* = .014) and matrices (incremental *R*^2^ = 1.47%, *p* = 005), were slightly weaker explained by CP-PGS (Figure 1). Interestingly, CP-PGS had a high predictive power for processing speed (incremental *R*^2^ = 2.66%, *p* = .001). The least predictive power was detected for individual differences in memory performance (incremental *R*^2^ = 1.07%, *p* = .014).

Individual differences in EA-PGS were especially predictive of individual differences in general intelligence (incremental *R*^2^ = 2.06%, *p* < .001). IST-2000-R verbal intelligence (incremental *R*^2^ = 1.61%, *p* = .004) and numerical intelligence (incremental *R*^2^ = 1.27%, *p* = .007) as well as aspects of general knowledge assessed by IST-2000-R (incremental *R*^2^ = 1.39%, *p* = .002) and BOWIT (incremental *R*^2^ = 1.25%, *p* = .002) were moderately predicted by EA-PGS. (Figure 1). Differences in language-free components of general intelligence, measured as figural intelligence (incremental *R*^2^ = 1.17%, *p* = .013) matrices (incremental *R*^2^ = 0.70%, *p* =.054) and processing speed (incremental *R*^2^ = 0.67%, *p* = .057) were partially only weakly explained by EA-PGS (Figure 1). Again, the predictive power of EA-PGS in individual differences in memory performance was weak (incremental *R*^2^ = 0.67%, *p* = .051) (Figure 1).

Apart from the *p*-value threshold (PT) of 0.05, we also investigated the predictive power of PGS with the strongest association with the respective cognitive tests (and subtests) in our sample, so-called best-fit PGS.

Here best-fit IQ-PGS were especially predictive of individual differences in general intelligence measured with the IST-2000-R (incremental *R*^2^ = 4.95%, *p* < .001). The predictive power for numerical intelligence and verbal intelligence was only slightly lower (Figure 2). Here, IQ-PGS had an incremental *R*^2^ of 4.41% for numerical intelligence (*p* < .001) and 4.20% for differences in verbal intelligence (*p* < .001). Regarding measures of general knowledge, IQ-PGS had an incremental *R*^2^ of 2.38 % for ability differences assessed with the IST-2000-R general knowledge test (*p* < .001) and 1.65% for differences in BOWIT (*p* < .001). Moreover, 2.00% of the difference in figural intelligence (*p* = .001), 2.04% of the performance difference in the BOMAT (*p* < .001) and 1.99% in processing speed (*p* < .001) were additionally explained by including IQ-PGS into the respective regression model. As depicted in Figure 2, IQ-PGS had a low predictive power for memory (incremental *R*^2^ = 1.08%, *p* = .013).

**Figure 2.**
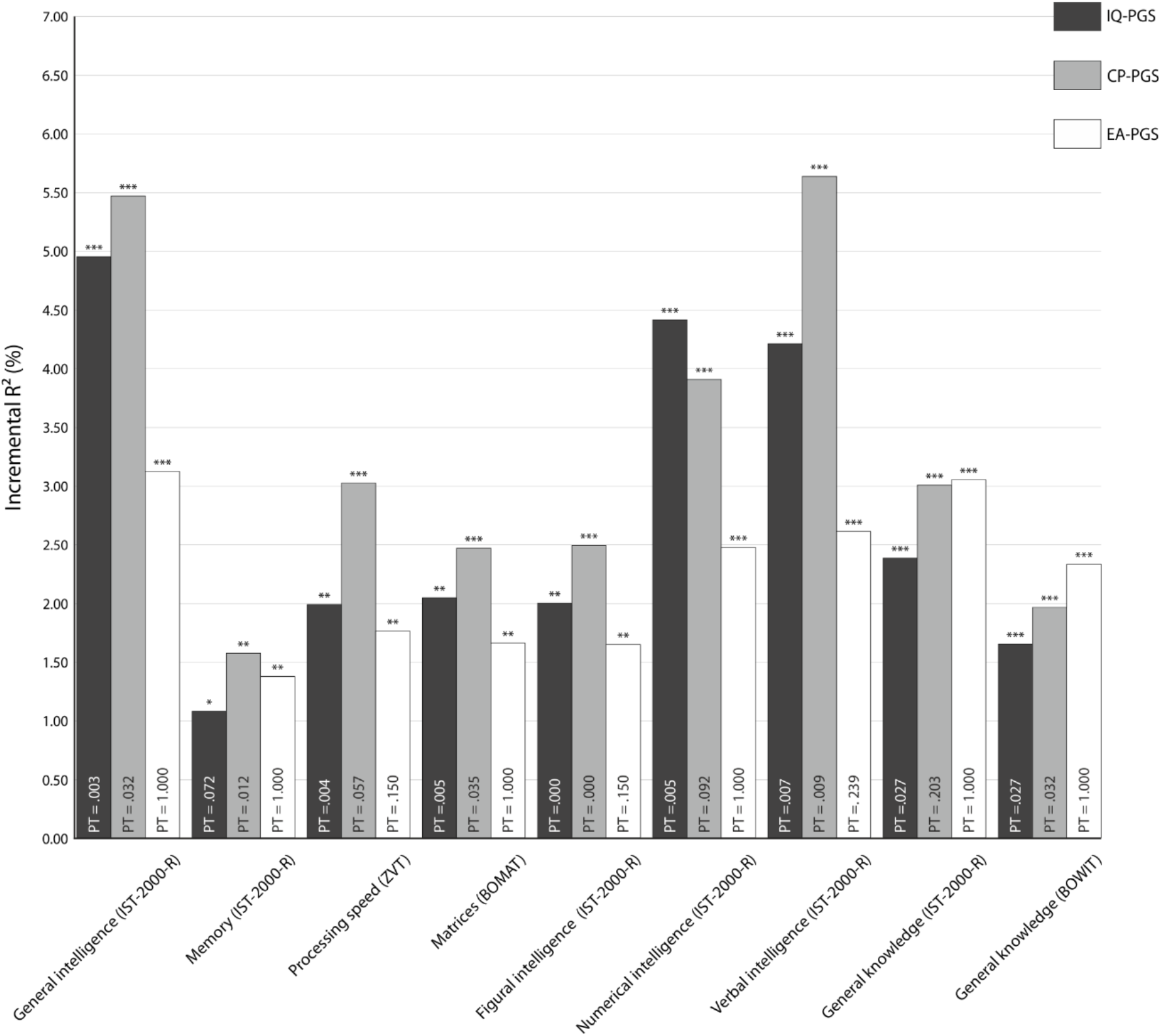
Incremental *R*^2^ of the best-fit polygenic scores of intelligence (IQ-PGS), cognitive performance (CP-PGS) and educational attainment (EA-PGS) in percent. The *p*-value thresholds (PT) that determined the inclusion of SNPs into the respective PGS are displayed in the respective bar. The incremental *R*^2^ reflects the increase in the determination coefficient (*R*^2^) when the IQ-PGS or CP-PGS or EA-PGS is added to a regression model predicting individual differences in the respective cognitive test. The association between PGS and phenotype was controlled for the effects of sex, age, population stratification, and multiple comparisons [40].* adjusted *p* ≤ .05, ** adjusted *p* ≤ .01, *** adjusted *p* ≤ .001.

Individual differences in CP-PGS were especially predictive of individual differences in verbal intelligence (incremental *R*^2^ = 5.63%, *p* < .001), general intelligence (incremental *R*^2^ = 5.47%, *p* < .001) and numerical intelligence (incremental *R*^2^ = 3.91%, *p* <.001) measured with the IST-2000-R (Figure 2). Aspects of general knowledge assessed by IST-2000-R (incremental *R*^2^ = 3.00%, *p* < .001) and BOWIT (incremental *R*^2^ = 1.96%, *p* < .001) were moderately predicted by CP-PGS. Differences in language-free components of general intelligence, measured as figural intelligence (incremental *R*^2^ = 2.49%, *p* < .001) and matrices (incremental *R*^2^ = 2.47%, *p* < .001), were also moderately explained by CP-PSG (Figure 2). Interestingly, CP-PGS had a higher predictive power for processing speed (incremental *R*^2^ = 3.02%, *p* < .001). The least predictive power was detected for individual differences in memory performance (incremental *R*^2^ = 1.57%, *p* = .003).

For EA-PGS we especially found a high predictive power for individual differences in general intelligence (incremental *R*^2^ = 3.12%, *p* < .001) and general knowledge assessed by the IST-2000-R (incremental *R*^2^ = 3.05%, *p* < .001). Verbal intelligence (incremental *R*^2^ = 2.61%, *p* < .001) and numerical intelligence (incremental *R*^2^ = 2.48%, *p* <. 001) measured with the IST-2000-R (Figure 2) as well as general knowledge assessed by BOWIT (incremental *R*^2^ = 2.33%, *p* < .001) were moderately predicted by EA-PGS. Differences in non-verbal aspects of intelligence were weaker predicted. Our multiple regression analyses resulted in predictive values of *R*^2^ = 1.76% for processing speed (*p* = .002), *R*^2^ = 1.66% for matrices (*p* = .003) and *R*^2^ = 1.64% for figural intelligence (*p* = .003). Again, the lowest predictive power was detected for individual differences in memory performance (incremental *R*^2^ = 1.38%, *p* = .005).

Next, we investigated the predictive power of PGS summarizing the effects of all SNPs (PT = 1.00, non-fit PGS, Figure 3). Here, again the increment in the determination coefficient caused by IQ-PGS was especially high for numerical (incremental *R*^2^ = 3.00%, *p* < .001), verbal (incremental *R*^2^ = 2.31%, *p* < .001), and general intelligence (incremental *R*^2^ = 1.94%, *p* = .001) measured with the IST-2000-R. Moreover, IQ-PGS increases the determination of the differences in general knowledge measured via IST-2000-R by 1.50% (*p* =.001) and by 1.21% (*p* =.002) for knowledge differences assessed by BOWIT. While differences in processing speed were predicted with an incremental *R*^2^ of 1.17% (*p* = .012), IQ-PGS had the lowest incremental effect on predicting differences in matrices (incremental *R*^2^ =0.60%, *p* = .074), memory (incremental *R*^2^ = 0.71%, *p* = .045) and figural intelligence (incremental *R*^2^ = 0.00%, *p* = .982).

**Figure 3.**
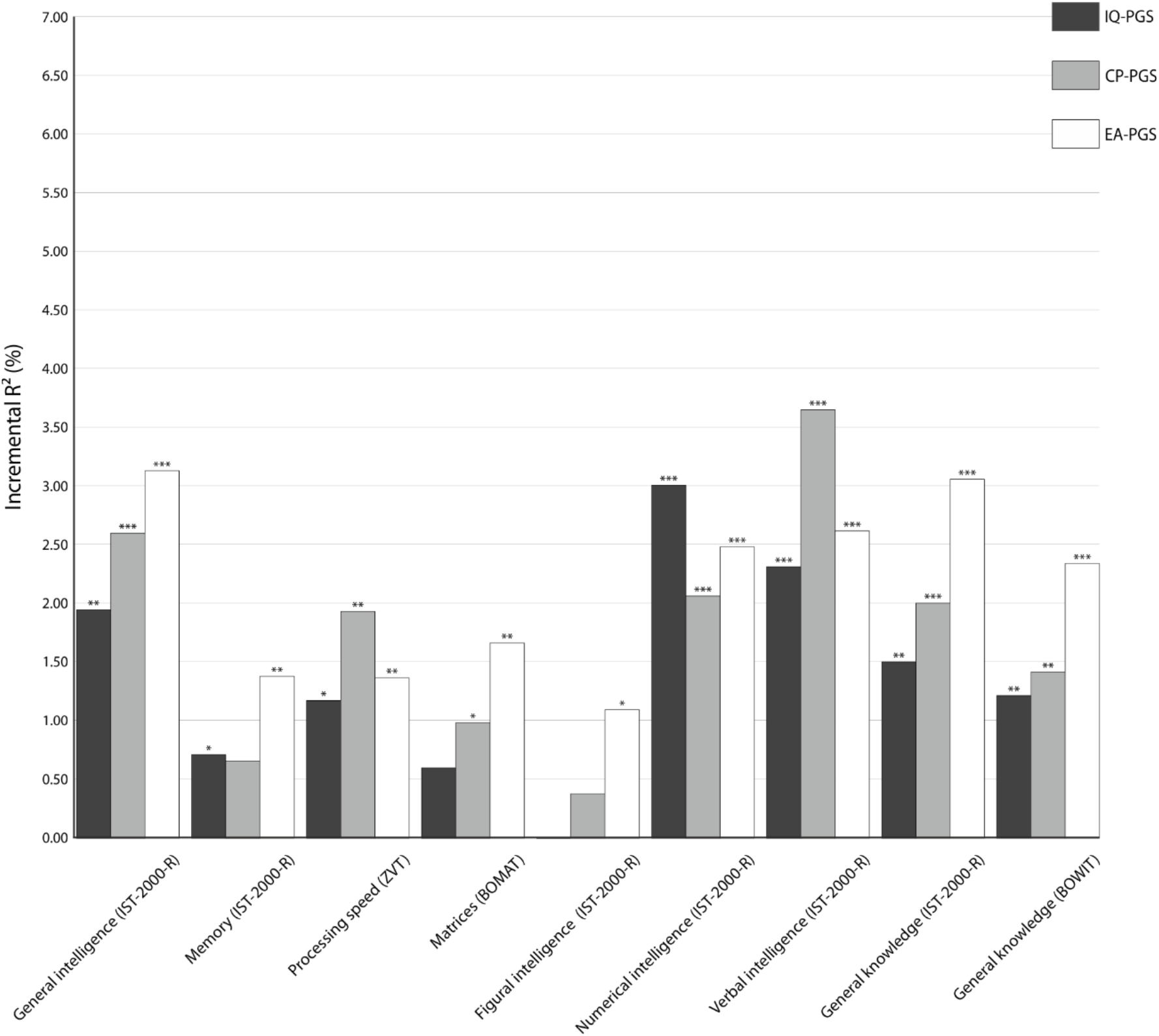
Incremental *R*^2^ of the non-fit polygenic scores of intelligence (IQ-PGS), cognitive performance (CP-PGS) and educational attainment (EA-PGS) in percent. P-value threshold (PT) = 1. The incremental *R*^2^ reflects the increase in the determination coefficient (*R*^2^) when the IQ-PGS or CP-PGS or EA-PGS is added to a regression model predicting individual differences in the respective cognitive test. The association between PGS and phenotype was controlled for the effects of age, sex, population stratification, and multiple comparisons [40]. *adjusted *p* ≤ .05, ** adjusted *p* ≤ .01, *** adjusted *p* ≤ .001.

A similar picture emerges for the predictive power of CP-PGS. The highest predictive power occurred for measures of verbal intelligence (incremental *R*^2^ = 3.64%, *p* < .001), general intelligence (incremental *R*^2^ = 2.59%, *p* < .001), numerical intelligence (incremental *R*^2^ = 2.06%, *p* < .001). The predictive power for differences in general knowledge assessed by IST-2000-R (incremental *R*^2^ = 1.99%, *p* <.001) and BOWIT (incremental *R*^2^ = 1.41%, *p* < .001) but also processing speed (incremental *R*^2^ = 1.93%, *p* = .001) were only slightly lower (Figure 3). Measures of non-verbal intelligence and memory were only poorly predicted by CP-PGS (; matrices: *R*^2^ = 0.99%, *p* =.022; figural intelligence: *R*^2^ = 0.38%, *p* = .158; memory: *R*^2^ = 0.65%, *p* =.054).

Individual differences in EA-PGS were especially predictive of individual differences in general intelligence (incremental *R*^2^ = 3.12%, *p* < .001), general knowledge assessed by IST-2000-R (incremental *R*^2^ = 3.05%, *p* < .001). General knowledge defined by BOWIT (incremental *R*^2^ = 2.33%, *p* < .001) as well as verbal (incremental *R*^2^ = 2.61%, *p* < .001) and numerical intelligence (incremental *R*^2^ = 2.48%, *p* < .001) were moderately predicted by EA-PGS. As shown in Figure 3, measures of non-verbal intelligence and memory were only poorly predicted by IQ-PGS (processing speed: *R*^2^ = 1.36%, *p* =.006; matrices: *R*^2^ = 1.65%, *p* = .004; figural intelligence: *R*^2^ = 1.09%, *p* =.017; memory: *R*^2^ = 1.38%, *p* =.005).

Overall, for the associations between cognitive measures and PT = 0.05, PT = 1.00 or best-fit PGS, a comparable pattern emerged, albeit the magnitude of explained variance deviated. Here, in general the predictive power of the PT = 0.05 and PT = 1.00 was somewhat lower than for the best-fit PGS. Moreover, the amount of explained variance was, in most cases, higher for IQ-PGS and CP-PGS than for EA-PGS (Figure 1, Figure 2, and Figure 3).

Finally, we tested whether there are sex-differences in the cognitive test scores and the predictive power of respective PGS. We did not observe a significant sex difference with regard to age (t(555) = 0.96, p = .304) and verbal intelligence (t(555) = 1.49, p = .14), processing speed (t(555) = .85, p = .39) and figural intelligence (t(555) = 1.32, p = .19). However, we found that females scored significantly higher on memory performance (t(555) = -4.87, p < .001), while males achieved significantly higher scores for general intelligence (t(555) = 5.01, p < .001), matrices (t(555) = 2.21, p = .03), numerical intelligence (t(555) = 7.65, p < .001), and general knowledge assessed by IST-2000-R (t(555) = 10.42, p < .001) and BOWIT (t(555) = 9.63, p < .001). Given the substantial sex differences for most of the cognitive test scores, we decided to compute the aforementioned predictive power of the respective PGS for both sexes separately (Figure S1 to S3). Although the pattern of results seems to be different for females and males, 78 out of 81 incremental *R*^2^ comparisons between the sexes were not significant (p >. 05). Statistical analysis indicated a significant sex-difference in incremental *R*^2^ of EA-PGS for matrices (p < .05) (PT = 0.05 EA-PGS; p = .04, best-fit EA-PGS; p = .0037, EA-PGS PT = 1.00; p = .0039) in the first place, however none of the p-values survived the control for multiple comparisons as defined by the Benjamini-Hochberg method.

## Discussion

Polygenic scores for intelligence, cognitive performance and educational attainment are increasingly used to investigate associations between genetic disposition for cognitive abilities and different life outcomes [21-23,25,24]. It is, however, currently not known which aspects of general intelligence are reflected to what extent by available PGS, derived from large GWAS. Here, we show that IQ-PGS, CP-PGS and EA-PGS do not predict every form of cognitive ability equally well (Figure 1 to 3). Specifically, we found for all three PGS for the whole group as well as separated for the sexes the pattern that all PGS had a high predictive power for interindividual differences in general, verbal and numerical intelligence. In contrast, memory was only weakly associated. The pattern for matrices was not consistent. For the whole group as well for males we found a poor association, whereas for females we found a higher predictive power.

Previous findings investigated the predictive power of single PGS in cognitive abilities. Liu et al. [41] used EA-PGS to predict verbal and matrix reasoning, and a recent preprint by Loughnan et al. [42] investigated the predictive power of IQ-PGS on different cognitive facets in children. Although, a direct comparison between these studies and our results is not straightforward since we report incremental *R2* for the predictive power and these studies used the standardized regression coefficient beta, by and large, our findings are in accordance with the results reported in these studies. More specifically, PGS estimated from cognitive ability approximations, like intelligence, cognitive performance, and educational attainment are more strongly associated with crystallized cognitive abilities compared to fluid abilities. This finding is counterintuitive at the first glance, since previous classical models [43] assumed that crystallized abilities would be less influenced by genetics but more impacted by environmental factors, like education. Nevertheless, recent evidence from a meta-analytic twin study showed higher heritability estimates for crystallized compared to fluid abilities [44]. Here, Kan et al. [44] speculated that these findings could be explained in terms of genotype-environment covariance. Because the acquisition of crystallized abilities (e.g. knowledge) depends on fluid abilities (e.g. cognitive processing) as described by the investment hypothesis [43], individuals who develop relatively high levels of cognitive-processing abilities tend to achieve relatively high levels of knowledge. More specifically, high achievers are more likely to end up in cognitively demanding environments which facilitate the initial genetic predisposition and thus the further development of a wide range of knowledge and skills. If, in addition, stimulating environments foster societally valued knowledge and skills more than cognitive processing per se, data simulations with dynamical models indicate that heritability coefficients of crystallized abilities could exceed those of fluid abilities [44]. Although this assumption seems plausible, we assume an alternative and more parsimonious explanation for these findings. We suggest that the predictive power of PGS in target samples could be influenced by the design of the GWAS that were used to discover the PGS in the first place. In typical GWAS, sample sizes tend to be very large, which usually comes at the cost of light phenotyping. It follows that PGS, which summarize trait-associated effect sizes of single SNPs, will reflect the genetic basis of the measured phenotype. In the case of educational attainment, this was defined as the years of schooling an individual completed [25]. Because verbal and numerical intelligence reflect culturally acquired abilities [2], it is not surprising, that PGS based on individual differences in years of education are primarily associated with individual differences in these aspects of intelligence, and less so with differences in non-verbal intelligence and memory. Of note, a recent analysis indicates that non-cognitive genetic factors, i.e. genetic variation in educational outcomes not explained by genetic variation in cognitive ability, accounted for more than half of the genetic variance in EA, and that heritable non-cognitive skills influence personality characteristics, and downstream health outcomes [45].

Interestingly, the same applies to the IQ-PGS and CP-PGS. These scores were based on a GWAS meta-analysis, that was mainly driven by the UK Biobank subsample. With almost 200,000 participants, the UK Biobank sample contributed at least above two-thirds of the meta-analysis sample [25]. In the UK Biobank, cognitive abilities were measured as the number of correct answers to a total of 13 questions assessing both verbal and mathematical intelligence. Although the respective questions are considered to measure intelligence in the form of verbal and numerical abilities, the number of correct answers seem to be dependent on culturally acquired knowledge rather than on deductive and inductive reasoning [46]. This becomes clear when taking a closer look at individual items. For example, arithmetic capability is measured with the following item: *“If David is twenty-one and Owen is nineteen and Daniel is nine years younger than David, what is half their age combined?” [46]*. As this is a classic word problem, it is not language-free and, therefore, clearly affected by cultural expertise. Again, the selection of test procedures used to determine individual differences in cognitive ability in the discovery sample appears to have a distorting influence on the predictive power of the PGS in other target samples. This is particularly important when one wants to investigate the functional pathways between genotype and phenotype. Researchers who aim to bridge the gap between genetic and neuronal correlates of cognitive abilities should be aware of this issue as previous studies have shown that distinct aspects of intelligence have distinct neural correlates [33]. Although our study provides important insights into the predictive power of commonly used PGS, the following limitations need to be considered. First, it has to be noted that with the present design, we cannot distinguish between direct genetic effects on cognitive performance and indirect, environmentally mediated parental genetic effects in form of genotype-environment correlations (e.g., cognitively enriching environments provided by parents [47]). Study designs combining family data with genotypic data, will be needed to further dissect those contributions [48]. Second, in the present study, the training data was primarily of European ancestry, and the target sample was a homogenous sample largely consisting of German university students. While polygenic prediction has been demonstrated to work best for discovery and target samples of matching ethnic background [49,42], as in the present study, the availability of large-scale discovery samples for other ancestries, e.g. of African or Asian populations, would be indispensable for the comparative analysis of polygenic contribution to cognition in different ancestries and cultures. Third, our sample was mainly composed of university students with a restricted age range. Previous studies show that our results are in accordance to associations patterns for various cognitive domains in children [42]. However, independent replication of our results in diverse samples with similar age ranges but different educational backgrounds is highly desirable [50].

Forth, the distinctive feature of our study is that we report the predictive power of various PGS separately for the sexes. Our results show that for both sexes, all three PGSs have higher predictive power for general, verbal and numerical intelligence lower for memory. The pattern for matrices was not consistent. For the whole group as well for males we found a poor association, whereas for females we found a very high predictive power especially for EA-PGS. However, one has to be careful with these interpretations, because the patterns of both sexes and the comparison do not differ significantly from each other when controlling for multiple comparisons. Since the analyses for both sexes are based on an exploratory analysis and were performed with significantly smaller sample sizes it is not surprising that they suffer from reduced statistical power compared to the whole group analyses. Therefore, we encourage future studies to systematically investigate sex differences in terms of the predictive power of various PGS in their cohorts.

Fifth, for the selection of a suitable *p*-value threshold for SNP inclusion, we were guided by the original publications of Savage et al. [24] and Lee et al. [25]. However, there is no clear specification as to which *p*-value threshold achieves the best overall predictive power regarding cognitive ability differences. Therefore, we additionally used a best-fit approach, which yielded different *p*-value thresholds for SNP inclusion for the different cognitive tests (Figure 2). Here, *p*-value thresholds for the various PGS ranged from PT = .0001 to PT = 1 but were highly intercorrelated (see Supplementary Material S3, S4, S5) so that the different best-fit scores are highly comparable regarding their SNP composition. As using a best-fit approach can lead to an overestimation of the observed explained variance [39], we also report the associations of non-fit PGS with a *p*-value threshold of PT = 1.00 (Figure 3). With this approach, PGS that take all available SNPs for either intelligence, cognitive performance or educational attainment into account were tested for associations with the different cognitive tests. Although PT = .05 and PT = 1.00 have a consistently lower predictive power than the best-fit PGS, they exhibit a comparable pattern of results. This confirms that polygenic scores of intelligence, cognitive performance and educational attainment derived from large GWAS [25,24] reflect some aspects of intelligence, e.g., verbal and numerical intelligence, more accurately than others, like memory. In conclusion, our study is the first which systematically investigates the association between polygenic scores of intelligence [24], cognitive performance and educational attainment [25] and a wide range of general intelligence aspects in healthy adults. The study shows how large-scale light-phenotyping GWAS studies with a strong statistical power to identify associated variants, and smaller comprehensive deep-phenotyping approaches with a fine-grained assessment of the phenotype of interest can complement each other. We demonstrate that the discovery GWAS for intelligence, cognitive performance and educational attainment do not reflect every form of cognitive ability equally well. Realizing that the way of phenotyping in large GWAS affects the predictive power of the resulting PGS [see also 51] is essential for all future studies planning to use PGS to unravel the genetic correlates of cognitive abilities in their samples.

## Supporting information

Cover Letter (Revision)

Response Letter (Revision)

Supplementary Material (Revision)

## Ethics Approval

The study protocol was approved by the local ethics committee of the Faculty of Psychology at Ruhr University Bochum (vote Nr. 165).

## Consent to Participate

All participants gave written informed consent and were treated following the Declaration of Helsinki.

## Consent for publication

All participants gave written informed consent that their data might be part of scientific publications. In case of publication, it is ensured that the published data cannot be associated with individual participants at any time.

## Availability of data and material

Since the data included in this study are part of a research project which is just completed, they are not openly accessible yet. However, if the manuscript is accepted for publication in Molecular Neurobiology, all necessary data will be made publicly available via an OSF link.

All cognitive ability scores were assessed via standardized and published test instruments. Publication of the corresponding material is therefore not permitted due to copyright regulations.

## Code availability

Analysis was applied with the freely available software PRSice 2.1.6. The specific code that was used for the analyses in this paper is available from the corresponding author upon reasonable request.

## Author Contributions

E.G., R.K. and S.O conceived the project and supervised the experiments. E.G., C.S., L.A., H.P.N, O.G., R.K, and S.O. designed the project. L.A., H.P.N., F.S., and R.K. planned and performed genetic experiments. C.S. and C.F. collected data. E.G., C.S., L.A., D.M., M.C.V., C.F., R.K., and S.O. analyzed the data. E.G., C.S., F.S., R.K. and S.O. wrote the paper. All authors discussed the results and edited the manuscript.

E.G. Erhan Genç

C.S. Caroline Schlüter

C.F. Christoph Fraenz

L.A. Larissa Arning

D.M. Dorothea Metzen

H.P.N Huu Phuc Nguyen

M.C.V Manuel C. Voelkle

F.S. Fabian Streit

O.G. Onur Güntürkün

R.K. Robert Kumsta

S.O. Sebastian Ocklenburg

## Acknowledgements

The authors would like to thank all research assistants for their support during the behavioral measurements.

## References

1. Neisser U, Boodoo G, Bouchard Jr TJ, Boykin AW, Brody N, Ceci SJ, Halpern DF, Loehlin JC, Perloff R, Sternberg RJ (1996) Intelligence: knowns and unknowns. Am Psychol 51 (2):77. doi:Doi 10.1037/0003-066x.51.2.77

2. Flanagan DP, Dixon SG (2013) The Cattell-Horn-Carroll Theory of Cognitive Abilities. Encyclopedia of Special Education: A Reference for the Education of Children, Adolescents, and Adults with Disabilities and Other Exceptional Individuals

3. Hossiep R, Hasella M, Turck D (2001) BOMAT-advanced-short version: Bochumer Matrizentest. Hogrefe, Göttingen, Germany

4. Raven JC (1990) Court, J.H., & Raven, J. (1990). Coloured progressive matrices. Manual for Raven’s Progressive Matrices and Vocabulary Scales

5. Hossiep R, Schulte M (2008) BOWIT: Bochumer Wissenstest. Hogrefe,

6. Roth B, Becker N, Romeyke S, Schafer S, Domnick F, Spinath FM (2015) Intelligence and school grades: A meta-analysis. Intelligence 53:118–137. doi:http://dx.doi.org/10.1016/j.intell.2015.09.002

7. Spengler M, Brunner M, Damian RI, Lüdtke O, Martin R, Roberts BW (2015) Student characteristics and behaviors at age 12 predict occupational success 40 years later over and above childhood IQ and parental socioeconomic status. Developmental psychology 51 (9):1329. doi:10.1037/dev0000025

8. Deary IJ, Whiteman MC, Starr JM, Whalley LJ, Fox HC (2004) The impact of childhood intelligence on later life: following up the Scottish mental surveys of 1932 and 1947. Journal of personality and social psychology 86 (1):130

9. Gottfredson LS, Deary IJ (2004) Intelligence predicts health and longevity, but why? Current Directions in Psychological Science 13 (1):1–4

10. Wraw C, Deary IJ, Der G, Gale CR (2016) Intelligence in youth and mental health at age 50. Intelligence 58:69–79

11. Knopik VS, Neiderhiser JM, DeFries JC, Plomin R (2016) Behavioral genetics. 7 edn. Macmillan Higher Education/W.H.Freeman & Co Ltd., London, UK

12. Plomin R, DeFries JC, Knopik VS, Neiderhiser JM (2016) Top 10 replicated findings from behavioral genetics. Perspectives on psychological science 11 (1):3–23

13. Fisher RA (1918) The correlation between relatives on the supposition of Mendelian inheritance. Earth and Environmental Science Transactions of the Royal Society of Edinburgh 52 (2):399–433

14. Plomin R, von Stumm S (2018) The new genetics of intelligence. Nature Reviews Genetics 19 (3):148

15. Wray NR, Lee SH, Mehta D, Vinkhuyzen AAE, Dudbridge F, Middeldorp CM (2014) Research Review: Polygenic methods and their application to psychiatric traits. J Child Psychol Psychiatry 55 (10):1068–1087. doi:10.1111/jcpp.12295

16. Davies G, Armstrong N, Bis JC, Bressler J, Chouraki V, Giddaluru S, Hofer E, Ibrahim-Verbaas CA, Kirin M, Lahti J (2015) Genetic contributions to variation in general cognitive function: a meta-analysis of genome-wide association studies in the CHARGE consortium (N= 53 949). Mol Psychiatr 20 (2):183–192

17. Sniekers S, Stringer S, Watanabe K, Jansen PR, Coleman JR, Krapohl E, Taskesen E, Hammerschlag AR, Okbay A, Zabaneh D (2017) Genome-wide association meta-analysis of 78,308 individuals identifies new loci and genes influencing human intelligence. Nature Genetics 49 (7):1107

18. Rietveld CA, Medland SE, Derringer J, Yang J, Esko T, Martin NW, Westra H-J, Shakhbazov K, Abdellaoui A, Agrawal A (2013) GWAS of 126,559 individuals identifies genetic variants associated with educational attainment. science 340 (6139):1467–1471

19. Okbay A, Beauchamp JP, Fontana MA, Lee JJ, Pers TH, Rietveld CA, Turley P, Chen G-B, Emilsson V, Meddens SFW (2016) Genome-wide association study identifies 74 loci associated with educational attainment. Nature 533 (7604):539

20. Okbay A, Wedow R, Kong E, Turley P, Lee J, Zacher M, Thom K, Nguyen AT, Maghzian O, Linner RK (2017) GWAS of educational attainment: phase 3-main results. Behavior Genetics 47 (6):699–700

21. Belsky DW, Moffitt TE, Corcoran DL, Domingue B, Harrington H, Hogan S, Houts R, Ramrakha S, Sugden K, Williams BS, Poulton R, Caspi A (2016) The Genetics of Success:How Single-Nucleotide Polymorphisms Associated With Educational Attainment Relate to Life-Course Development. Psychological science 27 (7):957–972. doi:10.1177/0956797616643070

22. Jansen PR, Polderman TJC, Bolhuis K, van der Ende J, Jaddoe VWV, Verhulst FC, White T, Posthuma D, Tiemeier H (2018) Polygenic scores for schizophrenia and educational attainment are associated with behavioural problems in early childhood in the general population. J Child Psychol Psychiatry 59 (1):39–47. doi:10.1111/jcpp.12759

23. Krapohl E, Patel H, Newhouse S, Curtis CJ, von Stumm S, Dale PS, Zabaneh D, Breen G, O’Reilly PF, Plomin R (2018) Multi-polygenic score approach to trait prediction. Mol Psychiatr 23 (5):1368–1374. doi:10.1038/mp.2017.163

24. Savage JE, Jansen PR, Stringer S, Watanabe K, Bryois J, De Leeuw CA, Nagel M, Awasthi S, Barr PB, Coleman JR (2018) Genome-wide association meta-analysis in 269,867 individuals identifies new genetic and functional links to intelligence. Nature genetics 50 (7):912

25. Lee JJ, Wedow R, Okbay A, Kong E, Maghzian O, Zacher M, Nguyen-Viet TA, Bowers P, Sidorenko J, Linnér RK (2018) Gene discovery and polygenic prediction from a genome-wide association study of educational attainment in 1.1 million individuals. Nature genetics 50 (8):1112

26. Cohen J (1992) A power primer. Psychological Bulletin 112 (1):155–159. doi:http://dx.doi.org/10.1037/0033-2909.112.1.155

27. Faul F, Erdfelder E, Buchner A, Lang A-G (2009) Statistical power analyses using G* Power 3.1: Tests for correlation and regression analyses. Behavior Research Methods 41 (4):1149 –1160. doi:https://doi.org/10.3758/BRM.41.4.1149

28. Liepmann D, Beauducel A, Brocke B, Amthauer R (2007) Intelligenz-Struktur-Test 2000 R (IST 2000 R). Manual (2. erweiterte und überarbeitete Aufl.). Göttingen: Hogrefe

29. Beauducel A, Brocke B, Liepmann D (2001) Perspectives on fluid and crystallized intelligence: facets for verbal, numerical, and figural intelligence. Pers Indiv Differ 30 (6):977–994. doi:Doi 10.1016/S0191-8869(00)00087-8

30. Oswald WD, Roth E (1987) Der Zahlen-Verbindungs-Test (ZVT). Hogrefe Verlag fuer Psychologie,

31. Oelhafen S, Nikolaidis A, Padovani T, Blaser D, Koenig T, Perrig WJ (2013) Increased parietal activity after training of interference control. Neuropsychologia 51 (13):2781–2790

32. Genç E, Fraenz C, Schlüter C, Friedrich P, Hossiep R, Voelkle MC, Ling JM, Güntürkün O, Jung RE (2018) Diffusion markers of dendritic density and arborization in gray matter predict differences in intelligence. Nature communications 9 (1):1905. doi:https://doi.org/10.1038/s41467-018-04268-8

33. Genç E, Fraenz C, Schlüter C, Friedrich P, Voelkle MC, Hossiep R, Güntürkün O (2019) The Neural Architecture of General Knowledge. European Journal of Personality 33 (5):589–605

34. Ackerman PL, Beier ME, Bowen KR (2002) What we really know about our abilities and our knowledge. Pers Indiv Differ 33 (4):587–605. doi:10.1016/s0191-8869(01)00174-x

35. Lynn R, Ivanec D, Zarevski P (2009) Sex Differences in General Knowledge Domains. Collegium Antropol 33 (2):515–520

36. Bratko D, Butkovic A, Chamorro-Premuzic T (2010) The genetics of general knowledge: A twin study from Croatia. Pers Indiv Differ 48 (4):403–407. doi:10.1016/j.paid.2009.11.006

37. Chang CC, Chow CC, Tellier LC, Vattikuti S, Purcell SM, Lee JJ (2015) Second-generation PLINK: rising to the challenge of larger and richer datasets. GigaScience 4 (1). doi:10.1186/s13742-015-0047-8

38. Purcell S, Chang C PLINK. www.cog-genomics.org/plink/1.9/.

39. Choi SW, O’Reilly PF (2019) PRSice-2: Polygenic Risk Score software for biobank-scale data. GigaScience 8 (7). doi:10.1093/gigascience/giz082

40. Benjamini Y, Hochberg Y (1995) Controlling the false discovery rate: a practical and powerful approach to multiple testing. Journal of the Royal statistical society: series B (Methodological) 57 (1):289–300

41. Liu M, Rea-Sandin G, Foerster J, Fritsche L, Brieger K, Clark C, Li K, Pandit A, Zajac G, Abecasis GR, Vrieze S (2020) Validating Online Measures of Cognitive Ability in Genes for Good, a Genetic Study of Health and Behavior. Assessment 27 (1):136–148. doi:10.1177/1073191117744048

42. Loughnan RJ, Palmer CE, Thompson WK, Dale AM, Jernigan TL, Fan CC (2021) Polygenic Score of Intelligence is More Predictive of Crystallized than Fluid Performance Among Children. bioRxiv:637512. doi:10.1101/637512

43. Cattell RB (1971) Abilities: Their structure, growth, and action.

44. Kan KJ, Wicherts JM, Dolan CV, van der Maas HL (2013) On the nature and nurture of intelligence and specific cognitive abilities: the more heritable, the more culture dependent. Psychological science 24 (12):2420–2428. doi:10.1177/0956797613493292

45. Demange PA, Malanchini M, Mallard TT, Biroli P, Cox SR, Grotzinger AD, Tucker-Drob EM, Abdellaoui A, Arseneault L, van Bergen E, Boomsma DI, Caspi A, Corcoran DL, Domingue BW, Harris KM, Ip HF, Mitchell C, Moffitt TE, Poulton R, Prinz JA, Sugden K, Wertz J, Williams BS, de Zeeuw EL, Belsky DW, Harden KP, Nivard MG (2021) Investigating the genetic architecture of noncognitive skills using GWAS-by-subtraction. Nature Genetics 53 (1):35–44. doi:10.1038/s41588-020-00754-2

46. Sudlow C, Gallacher J, Allen N, Beral V, Burton P, Danesh J, Downey P, Elliott P, Green J, Landray M (2015) UK biobank: an open access resource for identifying the causes of a wide range of complex diseases of middle and old age. PLoS medicine 12 (3):e1001779

47. Selzam S, Ritchie SJ, Pingault J-B, Reynolds CA, O’Reilly PF, Plomin R (2019) Comparing Within-and Between-Family Polygenic Score Prediction. The American Journal of Human Genetics 105 (2):351–363. doi:https://doi.org/10.1016/j.ajhg.2019.06.006

48. Bruins S, Dolan CV, Boomsma DI (2020) The Power to Detect Cultural Transmission in the Nuclear Twin Family Design With and Without Polygenic Risk Scores and in the Transmitted–Nontransmitted (Alleles) Design. Twin Research and Human Genetics 23 (5):265–270. doi:10.1017/thg.2020.76

49. Lewis CM, Vassos E (2020) Polygenic risk scores: from research tools to clinical instruments. Genome Medicine 12 (1):44. doi:10.1186/s13073-020-00742-5

50. Hanel PH, Vione KC (2016) Do student samples provide an accurate estimate of the general public? PloS one 11 (12):e0168354

51. Cai N, Revez JA, Adams MJ, Andlauer TFM, Breen G, Byrne EM, Clarke T-K, Forstner AJ, Grabe HJ, Hamilton SP, Levinson DF, Lewis CM, Lewis G, Martin NG, Milaneschi Y, Mors O, Müller-Myhsok B, Penninx BWJH, Perlis RH, Pistis G, Potash JB, Preisig M, Shi J, Smoller JW, Streit F, Tiemeier H, Uher R, Van der Auwera S, Viktorin A, Weissman MM, Kendler KS, Flint J, Consortium MDDWGotPG (2020) Minimal phenotyping yields genome-wide association signals of low specificity for major depression. Nature Genetics 52 (4):437–447. doi:10.1038/s41588-020-0594-5.

