## Supplementary material for "Polygenic Scores for Cognitive Abilities and their Association with Different Aspects of General Intelligence – a Deep Phenotyping Approach": Cover Letter (Revision)

LEIBNIZ RESEARCH CENTRE
FOR WORKING ENVIRONMENT
AND HUMAN FACTORS

Leibniz
Association

IfADo | Ardeystrasse 67 | D-44139 Dortmund | GERMANY

*Molecular Neurobiology*
Editorial Board

**Department of Psychology and Neurosciences**
Ardeystrasse 67
D-44139 Dortmund - Germany

**Dr. Erhan Genç**
Group leader Neuroimaging and Interindividual Differences

**www.ifado.de**

26.02.2021

**Cover Letter**

Dear Prof. Benedict C. Albensi,

please find enclosed the revised version of our manuscript entitled "*Polygenic Scores for Cognitive Abilities and their Association with Different Aspects of General Intelligence – a Deep Phenotyping Approach*" authored by Erhan Genç, Caroline Schlüter, Christoph Fraenz, Larissa Arning, Dorothea Metzen, Huu Phuc Nguyen, Manuel C. Voelkle, Fabian Streit, Onur Güntürkün, Robert Kumsta, and Sebastian Ocklenburg.

First of all, we would like to thank you and the reviewers for appreciating the value of our study and for the possibility to resubmit a revised version of our manuscript. Beyond that, we are pleased to report that the data collection and analysis of our project has been completed meantime, allowing us to present the results of our new sample with more than 500 individuals. The fact that the result pattern, by and large, remained the same for the final sample, underlines the robustness of our previous results. In addition, the increase in predictive power that came along with 200 additional subjects allowed us to split our datasets and compute separate analyses for male and female individuals. Thus, we were able to get exploratory insights in sex-differences in the predictive power of PGS.

Moreover, we addressed the effects of influencing factors, like parental and societal characteristics, and ethnic heritage in our discussion. Finally, both reviewers had some minor remarks, which we addressed in detail in the 'Response to Reviewers' document. Apart from that, both reviewers considered this work to be of high quality and of significance for the readership of *Molecular Neurobiology*.

Leibniz Research Centre for Working Environment and Human Factors
Scientific Directors: Prof. Dr. Jan G. Hengstler, Prof. Dr. Michael Nitsche, Prof. Dr. Edmund Wascher, Prof. Dr. Carsten Watzl
Commercial Director: Brigitta Thiel | Legal Entity: Forschungsgesellschaft für Arbeitsphysiologie und Arbeitsschutz e. V.
Bank: Helaba | IBAN: DE92 3005 0000 0001 1164 41 | BIC: WELADEDD
VAT # 314 5702 0136 DE 124651626 | VR 2255 Register of Association Dortmund

We would like to thank you and the reviewers for the valuable comments, which have substantially improved the manuscript. We hope that our response addresses the issues raised satisfactorily and that the manuscript is now suitable for publication.

We thank you for your editorial consideration.

Yours sincerely,

Dr. Erhan Genç
