## Supplementary material for "Polygenic Scores for Cognitive Abilities and their Association with Different Aspects of General Intelligence – a Deep Phenotyping Approach": Response Letter (Revision)

**Response to Reviews:**

**Response to the Editor and General Remarks**

First, we would like to thank the Editor for the possibility to resubmit a revised version of our manuscript and the reviewers for appreciating the value of our work. Their helpful comments have substantially improved the manuscript. Since the data collection and analysis of our project has been completed in the meantime, we are pleased to report the results of our new sample with more than 500 individuals. We think that the fact that the result pattern, by and large, remained the same for the final sample, speaks for the robustness of the previous results. In addition, the predictive power of the PGS substantially increased and with the addition of more than 200 new subjects to our samples, we felt confident to split our datasets and compute separate analyses for male and female individuals. Furthermore, we add some discussion about the findings with regard to crystallized to fluid abilities. Finally, we also decided to report the results for the PGS for cognitive performance (CP-PGS). The predictive power of CP-PGS was first introduced in the study by Lee et al. [1]. We considered it to be important to investigate its predictive power for different facets of general intelligence alongside IQ-PGS and EA-PGS. We therefore think our latest results provide even richer and more far-reaching insights that could help other researchers to better target their analyses in their cohorts. We also revised our manuscript according to the reviewers' recommendations. We hope that our response addresses the issues raised satisfactorily.

**Response to Reviewers**

Reviewer #2: This manuscript presents a study to investigate the association between polygenic scores of educational attainment and intelligence and different aspects of general intelligence in healthy adults. Authors found that PGS derived from light phenotyping GWAS do not reflect different facets of intelligence equally well, and thus should not be interpreted as genetic indicators of intelligence per se. The findings refine that how PGS are related to other traits or life outcomes.

Response: We would like to thank the reviewer for appreciating the value of our work and for the helpful remarks. We hope that our response addresses the issues raised satisfactorily and that the manuscript is now suitable for publication.

1. A note of caution should be added when interpreting the I-PGS: it should not simply be thought of as a proxy for genetics. Each individual in each study inherited half of their genome from each parent and so these genetic effects can also have indirect influences on their cognitive performance through the cognitively enriching environments that parents provide. This need to be discussed.

Response: We thank the reviewer for this important suggestion. This topic is now addressed in the discussion using the following section:

First, it has to be noted that with the present design, we cannot distinguish between direct genetic effects on cognitive performance and indirect, environmentally mediated parental genetic effects in form of genotype-environment correlations (e.g., cognitively enriching environments provided by parents [47]). Study designs combining family data with genotypic data, will be needed to further dissect those contributions [48].

2. In the current study PGS dataset was trained only on individuals of European ethnic ancestry. Training and validating PGS on different ancestry groups can cause issues. These issues can result from alleles being present for a certain ancestry group but being not represented in the mono-ethnic discovery GWAS. This needs to be discussed.

Response: We thank the reviewer for pointing out this matter in our manuscript. In the present study, the training data was primarily of European ancestry, and the target sample was a homogenous sample largely consisting of German university students. While polygenic prediction has been demonstrated to work best for discovery and target samples of matching ethnic backgrounds [2] as in the present study, the availability of large scale discoveries samples for other ancestries, e.g. of African or Asian populations, would be of high importance for the comparative analysis of polygenic contribution to cognition in different ancestries and cultures. To address this issue, we have revised our discussion:

Second, in the present study, the training data was primarily of European ancestry, and the target sample was a homogenous sample largely consisting of German university students. While polygenic prediction has been demonstrated to work best for discovery and target samples of matching ethnic background [28,29], as in the present study, the availability of large-scale discovery samples for other ancestries, e.g. of African or Asian populations, would be indispensable for the comparative analysis of polygenic contribution to cognition in different ancestries and cultures.

3. Authors did not mention that there is a sex-differences in their results. This need to be discussed. It would be interesting to present your data in a sex-dependent manner.

Response: We thank the reviewer for this important remark. In fact, we found sex differences for some cognitive test scores and decided to report the predictive power of respective PGS for females and males, separately in the revised manuscript. Although the pattern of incremental $R^2$ for most of the bars seems to be different (Figure S1 to Figure S3), none of the incremental $R^2$ comparisons between females and males were significantly different when controlling for multiple comparisons as defined by the Benjamini-Hochberg method. Since the analyses for both sexes were performed with significantly smaller sample sizes it is not surprising that they suffer from reduced statistical power compared to the whole group analyses. We address these points in the discussion and encourage future studies to systematically investigate sex differences in terms of the predictive power of various PGS in their cohorts.

4. Authors need to explain what was their rational for measuring the predictive value of PGS-IQ for short-term memory but not intermediate/long-term memory?

Response:

We thank the reviewer for bringing this inaccuracy to our attention. In our study, we measured memory. The term "short-term memory" is, therefore, misleading. We apologize for this mistake and now refer to it as “memory” throughout the manuscript.

5. Clarify the type of statistical analysis in all figure legends?

Response: We apologize that the statistical analysis in figure legends was apparently unclear. However, each figure legend (Manuscript and Supplements) contains the following sentence describing the respective statistics:

The incremental $R^2$ reflects the increase in the determination coefficient ($R^2$) when the IQ-PGS or CP-PGS or EA-PGS is added to a regression model predicting individual differences in the respective cognitive test. The association between PGS and phenotype was controlled for the effects of sex, age, population stratification, and multiple comparisons.

We hope that this meets your expectations and adequately describes the statistical analysis.

6. Figure title should explain the results.

Response: Thank you for this helpful suggestion. We have now provided a title to all figures, explaining the depicted results.

7. Provide definition of GWAS in the abstract.

Response: In accordance with the editor's suggestion, we added a definition of GWAS in the abstract.

8. Be consistent through the manuscript. Either r2 or R2?

Response: We thank the reviewer for pointing out this inconsistency in our manuscript. We revised our manuscript accordingly, and opted for $R^{2.}$

9. Remove the space between line 375-376 and 395-396.

Response: The spaces have been removed in accordance with the recommendation.

Reviewer #4: This is a lovely piece of work and I congratulate the authors for coming up with this kind of study. It is so interesting that I read it in one go without any difficulty in understanding the message and flow of the manuscript. The only thing that I feel should be added is a strong real time message for the use of the findings of this study in future works.

Response: We would like to thank reviewer 4 for appreciating the value of our work. In our opinion, the real time massage of our work will be explained in the following sections:

[Introduction] This has been successfully achieved using a light-phenotyping approach, that is, performing GWAS on the performance in rather superficial tests of general cognitive abilities [24,25], or even more crudely, years of education [25]. The question thus arises, which of the various aspects of general intelligence [2,1], are mainly reflected in those GWAS. The study at hand aimed to tackle this issue by pursuing a deep phenotyping approach. Using an extensive test battery comprised of tests for memory performance, processing speed, reasoning, and general knowledge, we investigated the predictive power of IQ-PGS [24], CP-PGS and EA-PGS [25] with regard to each of the aforementioned cognitive abilities.

[Discussion] In conclusion, our study is the first which systematically investigates the association between polygenic scores of intelligence [24], cognitive performance and educational attainment [25] and a wide range of general intelligence aspects in healthy adults. The study shows how large scale light-phenotyping GWAS studies with a strong statistical power to identify associated variants, and smaller comprehensive deep-phenotyping approaches with a fine-grained assessment of the phenotype of interest can complement each other. We demonstrate that the discovery GWAS for intelligence, cognitive performance and educational attainment do not reflect every form of cognitive ability equally well. Realizing that the way of phenotyping in large GWAS affects the predictive power of the resulting PGS [see also 52] is essential for all future studies planning to use PGS to unravel the genetic correlates of cognitive abilities in their samples.

However, if reviewer 4 has a different understanding of strong real time massage, we would be pleased to receive further suggestions on this matter.

References

1. Lee JJ, Wedow R, Okbay A, Kong E, Maghzian O, Zacher M, Nguyen-Viet TA, Bowers P, Sidorenko J, Linnér RK (2018) Gene discovery and polygenic prediction from a genome-wide association study of educational attainment in 1.1 million individuals. Nature genetics 50 (8):1112

2. Lewis CM, Vassos E (2020) Polygenic risk scores: from research tools to clinical instruments. Genome Medicine 12 (1):44. doi:10.1186/s13073-020-00742-5
