## Supplementary Material (Revision) for "Polygenic Scores for Cognitive Abilities and their Association with Different Aspects of General Intelligence – a Deep Phenotyping Approach"

Genç et al. (2021)

### Supplemental Material

#### Table S1 Descriptive Statistics for all Cognitive measures.

| | Min | Max | *M* | *SD* |
|---|---|---|---|---|
| **General intelligence** (IST-2000-R) | 44.00 | 164.00 | 118.83 | 19.67 |
| **Memory** (IST-2000-R) | 3.00 | 23.00 | 17.07 | 4.06 |
| **Processing speed** (ZVT) | 32.00 | 95.25 | 54.59 | 10.51 |
| **Matrices** (BOMAT) | 3.00 | 28.00 | 16.02 | 4.13 |
| **Figural intelligence** (IST-2000-R) | 12.00 | 55.00 | 35.38 | 8.03 |
| **Numerical intelligence** (IST-2000-R) | 12.00 | 60.00 | 43.14 | 10.56 |
| **Verbal intelligence** (IST-2000-R) | 15.00 | 55.00 | 40.31 | 6.15 |
| **General knowledge** (IST-2000-R) | 15.00 | 80.00 | 53.31 | 9.79 |
| **General knowledge** (BOWIT) | 61.00 | 295.00 | 145.76 | 39.03 |

**Note.** Table S1 shows descriptive statistics for all cognitive measure raw test scores. Min = minimum test score, Max = maximum test score, *M* = mean test score and *SD* = standard deviation of test score. *N* = 557.

**Table S2 Intercorrelations between all Raw Test Scores for Cognitive Performance**

| | General intelligence (IST-2000-R) | Memory (IST-2000-R) | Processing speed (ZVT) | Matrices (BOMAT) | Figural intelligence (IST-2000-R) | Numerical intelligence (IST-2000-R) | Verbal intelligence (IST-2000-R) | General knowledge (IST-2000-R) |
|---|---|---|---|---|---|---|---|---|
| **Memory** (IST-2000-R) | .395*** | | | | | | | |
| **Processing speed** (ZVT) | -.571*** | -.419*** | | | | | | |
| **Matrices** (BOMAT) | .607*** | .442*** | -.420*** | | | | | |
| **Figural intelligence** (IST-2000-R) | .784*** | .338*** | -.430*** | .522*** | | | | |
| **Numerical intelligence** (IST-2000-R) | .860*** | .312*** | -.545*** | .521*** | .471*** | | | |
| **Verbal intelligence** (IST-2000-R) | .700*** | .287*** | -.332*** | .367*** | .396*** | .422*** | | |
| **General knowledge** (IST-2000-R) | .594*** | .162** | -.214*** | .331*** | .356*** | .537*** | .514*** | |
| **General knowledge** (BOWIT) | .408*** | .060 | -.051 | .168*** | .183*** | .359*** | .452*** | .834*** |

**Note.** Table S2 shows the Pearson's correlation coefficients for the intercorrelations between the test scores of all cognitive measures. $N$ = 557, *** $p \leq .001$, ** $p \leq .01$, * $p \leq .05$.

### Table S3 Intercorrelations between best-fit IQ-PGS

| Polygenic score best-fit for… | General intelligence (IST-2000-R) PT = .0029 | Memory (IST-2000-R) PT = .0719 | Processing speed (ZVT) PT = .0037 | Matrices (BOMAT) PT = .0055 | Figural intelligence (IST-2000-R) PT = .0003 | Numerical intelligence (IST-2000-R) PT = .0055 | Verbal intelligence (IST-2000-R) PT = .0071 | General knowledge (IST-2000-R) PT = .0266 |
|---|---|---|---|---|---|---|---|---|
| **Memory** (IST-2000-R) PT = .0719 | .702*** | | | | | | | |
| **Processing speed** (ZVT) PT = .0037 | .997*** | .727*** | | | | | | |
| **Matrices** (BOMAT) PT = .0055 | .934*** | .753*** | .962*** | | | | | |
| **Figural intelligence** (IST-2000-R) PT = .0003 | .814*** | .553*** | .790*** | .755*** | | | | |
| **Numerical intelligence** (IST-2000-R) PT = .0055 | .934*** | .753*** | .962*** | 1.000*** | .755*** | | | |
| **Verbal intelligence** (IST-2000-R) PT = .0071 | .908*** | .782*** | .938*** | .973*** | .744*** | .973*** | | |
| **General knowledge** (IST-2000-R) PT = .0266 | .786*** | .912*** | .810*** | .845*** | .635*** | .845*** | .875*** | |
| **General knowledge** (BOWIT) PT = .0266 | .786*** | .912*** | .810*** | .845*** | .635*** | .845*** | .875*** | 1.000*** |

**Note.** To estimate the comparability of the best-fit polygenic scores (PGS) correlation coefficients between each of the scores were calculated. Table S3 shows the Pearson's correlation coefficient for each PGS pair. PT = *p*-value threshold. *N* = 518, *** *p* ≤ .001, ** *p* ≤ .01, * *p* ≤ .05.

### Table S4 Intercorrelations between best-fit CP-PGS

| Polygenic score best-fit for… | General intelligence (IST-2000-R) PT = .0317 | Memory (IST-2000-R) PT = .0119 | Processing speed (ZVT) PT = .0570 | Matrices (BOMAT) PT = .0350 | Figural intelligence (IST-2000-R) PT = .0001 | Numerical intelligence (IST-2000-R) PT = .0918 | Verbal intelligence (IST-2000-R) PT = .0089 | General knowledge (IST-2000-R) PT = .2030 |
|---|---|---|---|---|---|---|---|---|
| **Memory** (IST-2000-R) PT = .0119 | .915*** | | | | | | | |
| **Processing speed** (ZVT) PT = .0570 | .948*** | .861*** | | | | | | |
| **Matrices** (BOMAT) PT = .0350 | .992*** | .906*** | .960*** | | | | | |
| **Figural intelligence** (IST-2000-R) PT = .0001 | .581*** | .644*** | .539*** | .567*** | | | | |
| **Numerical intelligence** (IST-2000-R) PT = .0918 | .914*** | .837*** | .958*** | .922*** | .517*** | | | |
| **Verbal intelligence** (IST-2000-R) PT = .0089 | .889*** | .973*** | .830*** | .879*** | .676*** | .805*** | | |
| **General knowledge** (IST-2000-R) PT = .2030 | .842*** | .775*** | .885*** | .850*** | .472*** | .935*** | .751*** | |
| **General knowledge** (BOWIT) PT = .0323 | .999*** | .914*** | .950*** | .993*** | .578*** | .916*** | .888*** | .846*** |

**Note.** To estimate the comparability of the best-fit polygenic scores (PGS) correlation coefficients between each of the scores were calculated. Table S4 shows the Pearson's correlation coefficient for each PGS pair. PT = *p*-value threshold. $N$ = 518, $^{***}$ $p \leq .001$, $^{**}$ $p \leq .01$, $^{*}$ $p \leq .05$

**Table S5 Intercorrelations between best-fit EA-PGS**

| Polygenic score best-fit for… | General intelligence (IST-2000-R) PT = 1 | Memory (IST-2000-R) PT = 1 | Processing speed (ZVT) PT = .1500 | Matrices (BOMAT) PT = 1 | Figural intelligence (IST-2000-R) PT = .1500 | Numerical intelligence (IST-2000-R) PT = 1 | Verbal intelligence (IST-2000-R) PT = .2390 | General knowledge (IST-2000-R) PT = 1 |
|---|---|---|---|---|---|---|---|---|
| **Memory** (IST-2000-R) PT = 1 | 1.000$^{***}$ | | | | | | | |
| **Processing speed** (ZVT) PT = .1500 | .947$^{***}$ | .947$^{***}$ | | | | | | |
| **Matrices** (BOMAT) PT = 1 | 1.000$^{***}$ | 1.000$^{***}$ | .947$^{***}$ | | | | | |
| **Figural intelligence** (IST-2000-R) PT = .1500 | 1.000$^{***}$ | 1.000$^{***}$ | .947$^{***}$ | 1.000$^{***}$ | | | | |
| **Numerical intelligence** (IST-2000-R) PT = 1 | .968$^{***}$ | .968$^{***}$ | .978$^{***}$ | .968$^{***}$ | .968$^{***}$ | | | |
| **Verbal intelligence** (IST-2000-R) PT = .2390 | 1.000$^{***}$ | 1.000$^{***}$ | .947$^{***}$ | 1.000$^{***}$ | 1.000$^{***}$ | .968$^{***}$ | | |
| **General knowledge** (IST-2000-R) PT = 1 | 1.000$^{***}$ | 1.000$^{***}$ | .947$^{***}$ | 1.000$^{***}$ | 1.000$^{***}$ | .968$^{***}$ | 1.000$^{***}$ | |
| **General knowledge** (BOWIT) PT = 1 | .947$^{***}$ | .947$^{***}$ | 1.000$^{***}$ | .947$^{***}$ | .947$^{***}$ | .978$^{***}$ | .947$^{***}$ | .947$^{***}$ |

**Note.** To estimate the comparability of the best-fit polygenic scores (PGS) correlation coefficients between each of the scores were calculated. Table S5 shows the Pearson's correlation coefficient for each PGS pair. PT = *p*-value threshold. $N$ = 518, $^{***}$ $p \leq .001$, $^{**}$ $p \leq .01$, $^{*}$ $p \leq .05$.

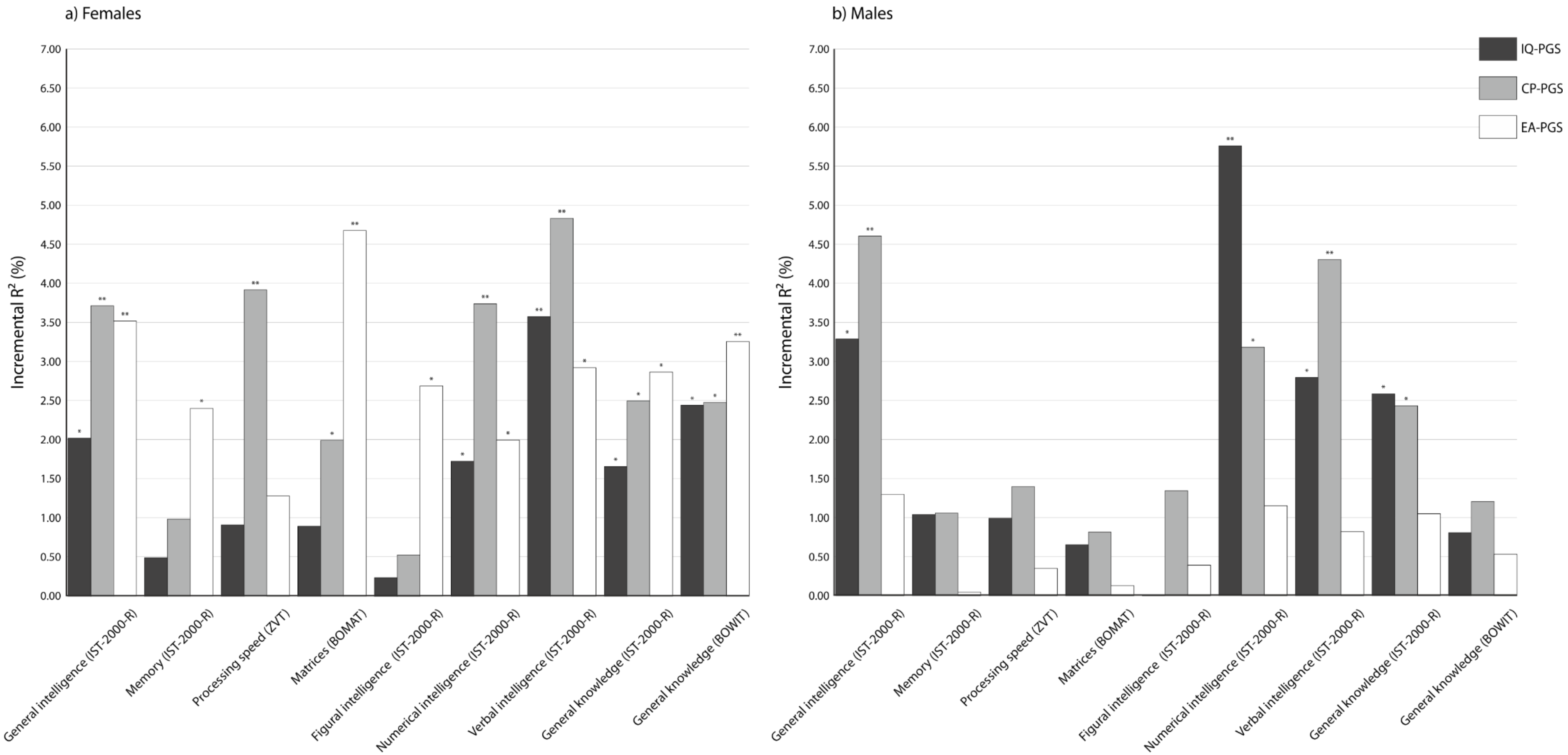


**Figure S1. Incremental $R^2$ of the *P*-value threshold (PT) = .05 polygenic scores of intelligence (IQ-PGS), cognitive performance (CP-PGS) and educational attainment (EA-PGS) in percent for females (a) and males (b)**. The incremental $R^2$ reflects the increase in the determination coefficient ($R^2$) when the IQ-PGS or CP-PGS or EA-PGS is added to a regression model predicting individual differences in the respective cognitive test. The association between PGS and phenotype was controlled for the effects of sex, population stratification, and multiple comparisons. $^{*}$ adjusted $p \leq .05$, $^{**}$ adjusted $p \leq .01$, $^{***}$ adjusted $p \leq .001$.

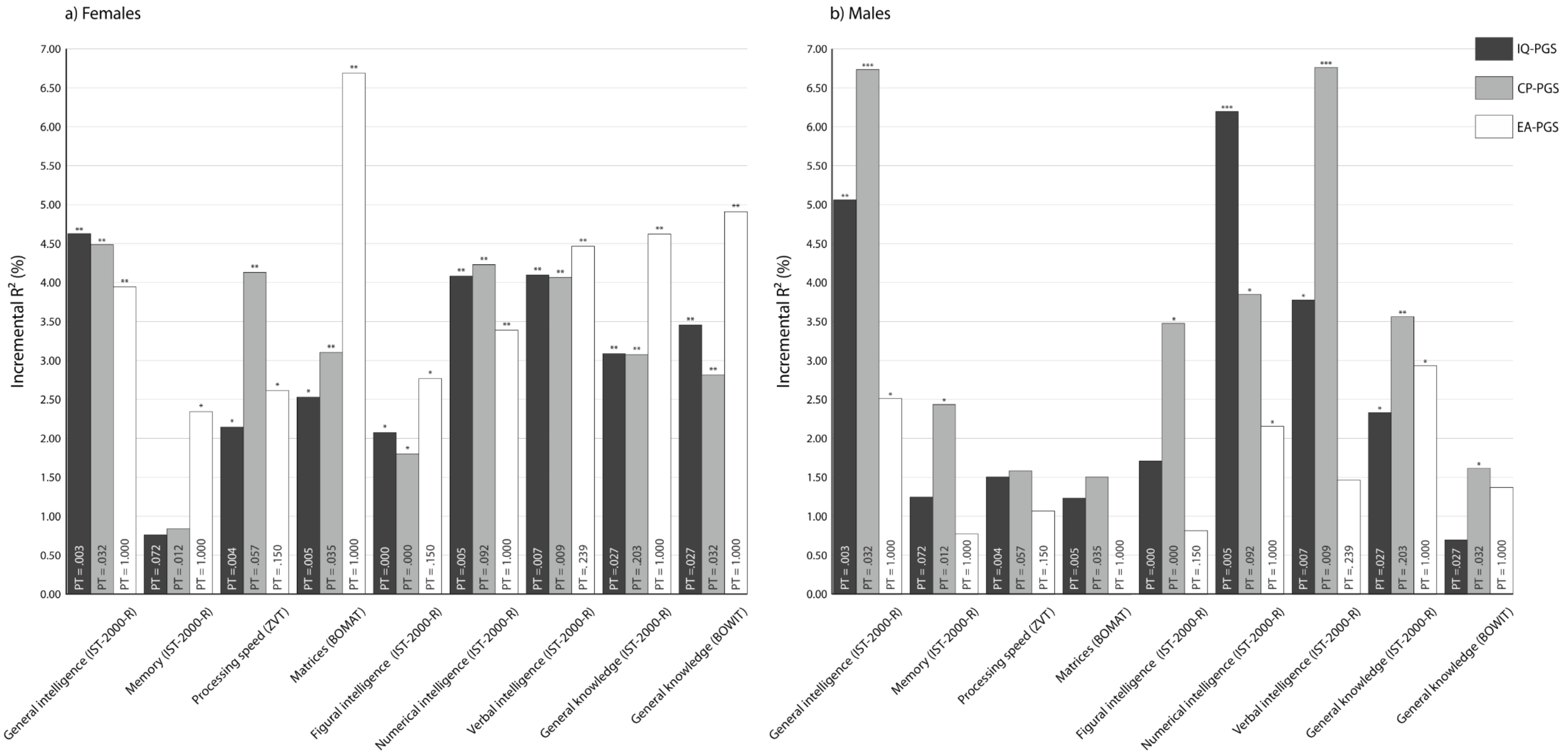


**Figure S2. Incremental $R^2$ of the best-fit polygenic scores of intelligence (IQ-PGS), cognitive performance (CP-PGS) and educational attainment (EA-PGS) in percent for females (a) and males (b)**. The *p*-value thresholds (PT) that determined the inclusion of SNPs into the respective PGS are displayed in the respective bar. The incremental $R^2$ reflects the increase in the determination coefficient ($R^2$) when the IQ-PGS or CP-PGS or EA-PGS is added to a regression model predicting individual differences in the respective cognitive test. The association between PGS and phenotype was controlled for the effects of age, population stratification, and multiple comparisons. $^{*}$ adjusted $p \leq .05$, $^{**}$ adjusted $p \leq .01$, $^{***}$ adjusted $p \leq .001$.

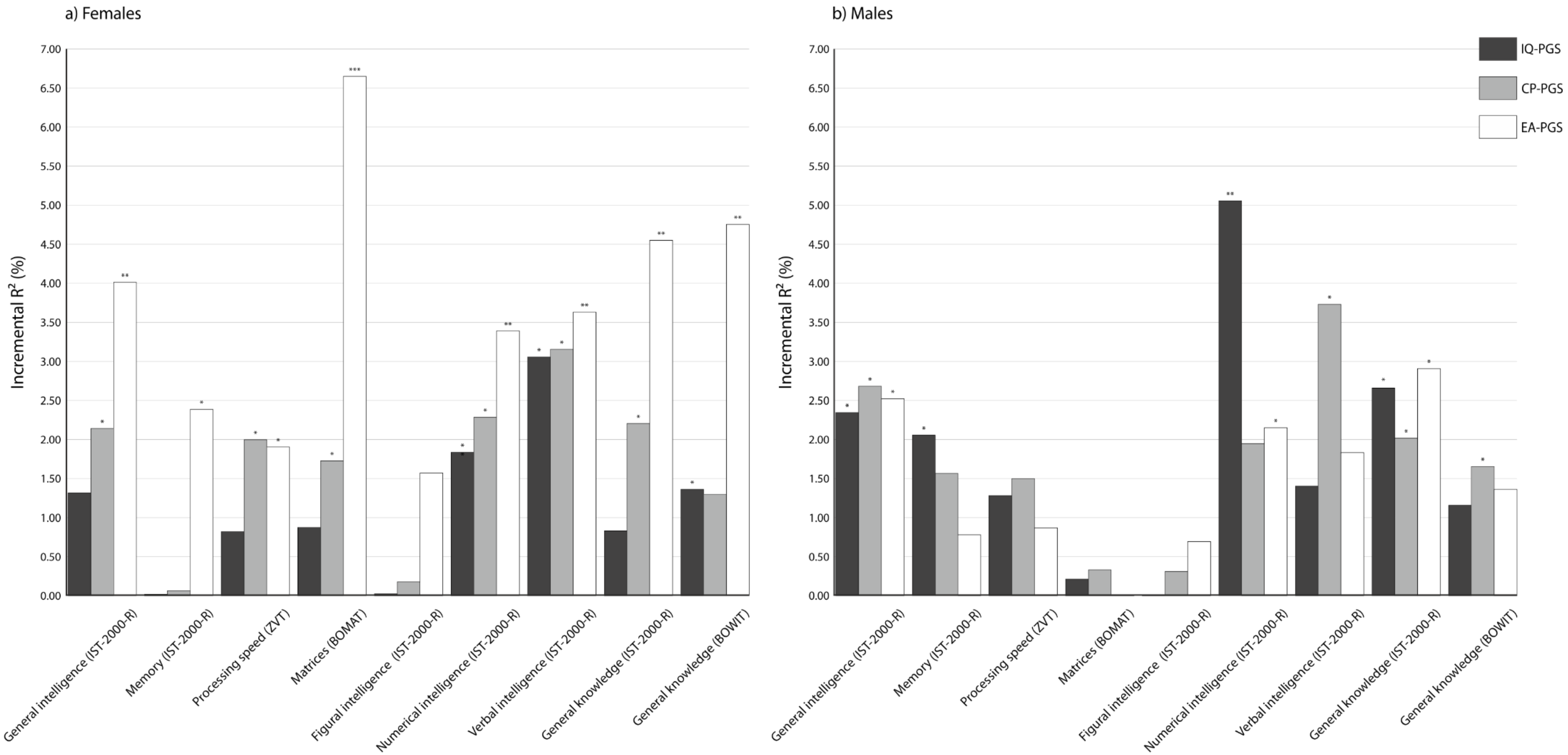


**Figure S3**. **Incremental** $R^2$ **of the non-fit (PT = 1) polygenic scores of intelligence (IQ-PGS), cognitive performance (CP-PGS) and educational attainment (EA-PGS) in percent for females (a) and males (b)**. The incremental $R^2$ reflects the increase in the determination coefficient ($R^2$) when the IQ-PGS or CP-PGS or EA-PGS is added to a regression model predicting individual differences in the respective cognitive test. The association between PGS and phenotype was controlled for the effects of age, population stratification, and multiple comparisons. $^{*}$ adjusted $p \leq .05$, $^{**}$ adjusted $p \leq .01$, $^{***}$ adjusted $p \leq .001$.
